# A Novel Method for Culturing Telencephalic Neurons in Axolotls

**DOI:** 10.1101/2025.02.14.638296

**Authors:** Sevginur Bostan, Safiye Serdengeçtİ, F. Kemal Bayat, Sadık Bay, Ayşe Server Sezer, Neşe Ayşİt, Gürkan Öztürk

## Abstract

The axolotl (*Ambystoma mexicanum*), a neotenic salamander with remarkable regenerative capabilities, serves as a key model for studying nervous system regeneration. Despite its potential, the cellular and molecular mechanisms underlying this regenerative capacity remain poorly understood, partly due to the lack of reliable *in vitro* models for axolotl neural cells. In this study, we developed a novel protocol for primary cultures of adult axolotl telencephalon/pallium, enabling the maintenance of viable and functionally active neural cells. Using calcium imaging and immunocytochemistry, we demonstrated the presence of neuronal and glial markers, synaptic connections, and spontaneous calcium activity, highlighting the functional integrity of the cultured cells. Our findings reveal that these cultures can be maintained in both serum and serum-free conditions, with neurons exhibiting robust neurite outgrowth and responsiveness to injury. This protocol addresses a critical gap in axolotl research by providing a controlled *in vitro* system to study neurogenesis and regeneration. By offering insights into the regenerative mechanisms of axolotl neurons, this work lays the foundation for comparative studies with mammalian systems, potentially informing therapeutic strategies for neurodegenerative diseases and CNS injuries in humans.

## Introduction

The Mexican salamander, axolotl (Ambystoma mexicanum), an urodele amphibian that cannot complete metamorphosis and preserves neotenic characteristics throughout its lifespan, has been a focus of interest as a model organism in regeneration research [1–3]. Compared to rather limited regeneration ability of nervous system tissues in adult mammals, urodele amphibians have a much higher capacity to regenerate their nervous system after an injury [4–7].

Telencephalon region in the brain of these amphibians is known to contain a highly proliferative and regenerative ventricular zone (VZ) compared to other reptiles [6]. There is very limited information on the structural and molecular mechanisms underlying this high regenerative capacity. The cellular organisation and characteristics of different cell populations of the axolotl brain and how they functionally regenerate after injury have not been well defined. This is partially due to lack of reliable *in vitro* models to study regenerative processes under controlled conditions of cell culture. In general, *in vitro* studies using axolotl cell cultures are scarce and existing ones mostly focus on blastema - derived cells [8]. Only available axolotl cell line is AL-1, whose origin is unclear [9]. Primary cultures of neural cells are even less common; the only example is a protocol for generation of neurospheres from newt brain [10, 11].

In this study, we developed a protocol for primary cultures of adult axolotl telencephalon/pallium. We tested the viability and maintenance of the primary culture with serum or serum-free medium. We also carried out calcium imaging to demonstrate they are functionally active and responsive. Additionally, we performed immunocytochemistry to determine cell types and synaptic connections in cultures.

## Materials and Methods

### 1. Ethics Statement and Animal Handling

Axolotls were kept and bred at Experimental Animal Centre of Istanbul Medipol University (MEDITAM). All animals were handled in strict accordance with guidelines for animal care and use issued by the EU directive code; 86/609/CEE. The Committee on Ethics of Animal Experimentation of Istanbul Medipol University (IMUHADYEK) approved all experimental procedures. Adult white axolotls (n=6), 12-16 cm in length, were used in the optimization of the culture protocol and experiments.

### 2. Primary Telencephalon Culture Protocol

#### Animal Surgery

Axolotls were anaesthetized with 0,02 % benzocaine (Sigma-Aldrich, E1501) and decapitated swiftly. The skull was cut from the spinal cord to the median fontanel, following a straight line. Upon reaching the median fontanel, a triangular section was excised, as depicted in Figure 1A. The telencephalon was accessed at the site where the skull piece was removed, as indicated in Figure 1A. Using a forceps with curved tips, the telencephalon was carefully isolated and transferred into the dissection medium Leibovitz’s L-15 (L-15-Gibco) (1% Pen-Strep (Sigma-Aldrich) and 1% L-glutamine (Sigma-Aldrich)). Under a stereo microscope, the meninges were removed, and extracted tissue is shown in Figure 1B.

**Figure 1.**
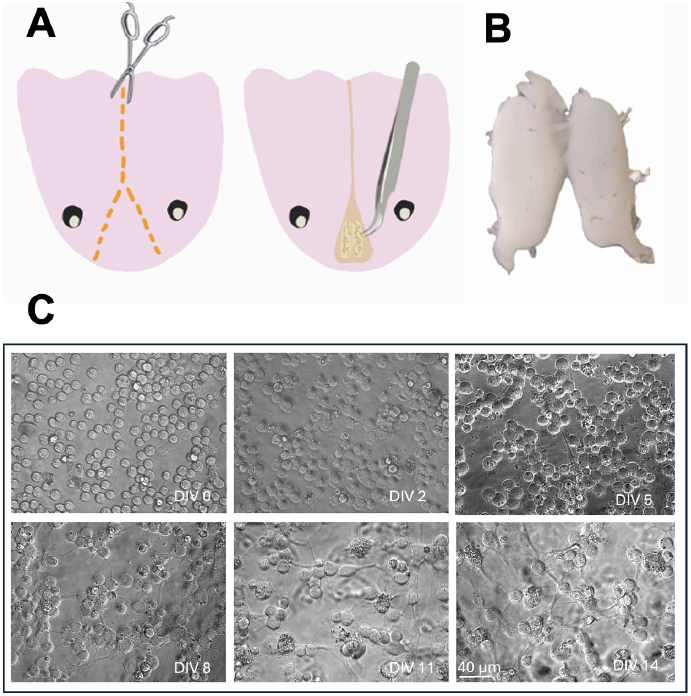
Axolotl primary telencephalon culture procedure **A** Dissection and removal of the brain **B** Isolated telencephalon **C** Brightfield images of cultured neurons during 2 weeks of incubation.

#### Tissue Dissociation

Culturing protocol was based on an earlier study [12] with several adjustments for axolotls. After dissection, both telencephalon hemispheres were transferred into a separate dissociation medium L-15 (1% Pen-Strep, 1% L-glutamine) with Papain (12.5 U/mL, Sigma-Aldrich) and incubated for 45 minutes at 4°C. Afterwards, DNAse (50 μg/mL, BioMatik) was added, and the tissue pieces were gently triturated with fire-polished glass Pasteur pipettes of decreasing diameters until a homogenous cell suspension was obtained. Then, 10% FBS (Sigma-F4135) is added to the cell suspension, and it is incubated for 15 minutes at room temperature for enzyme inhibition.

#### Gradient Centrifugation

To obtain a neuron–rich cell population free of tissue debris, a gradient centrifugation step was added, which was adapted from a previous study [13]. The cell suspension was carefully added on top of a four-layer Percoll (Sigma) gradient which was prepared in L-15 medium with 60, 30, 20 and 10% densities. Then it was spun at 3000 *xg* for 25 minutes in a centrifuge at 15 °C. The cells were collected from 30% layer, transferred into a fresh L-15 medium (1% Pen-Strep, 1% L-glutamine) and centrifuged for five minutes at 300 *xg* to remove Percoll.

#### Cell Seeding

After discarding the supernatant, cells were resuspended with the culture medium Neurobasal-A (NBA-Gibco) (1% Pen-Strep, 1% L-glutamine, 2% B-27 (Gibco), with or without 2% horse serum (Sigma, H1270). Prior to seeding, polystyrene cloning cylinders (I.D. × H 4.7 mm × 8 mm, Sigma, C7983) were placed on 35 mm diameter dishes (WPI) pre-coated with PEI (0.05 - 0.1%, Sigma). Then, the cells were seeded into the cloning cylinders at three different confluencies; High Confluency (HC) (12000/140 μl), Medium Confluency (MC) (6000/140 μl) and Low Confluency (LC) (3000/140 μl). Culture plates were maintained in an incubator at 21°C with 2% CO_2_. Thereafter, 30-40 μl of fresh medium was added every 3-4 days. The cloning cylinders were removed on DIV14, and 1 mL of fresh culture medium was added to the plates, and half of the medium was replaced every 3-4 days. To test viability and observe the development of neurons *in vitro*, cultures were maintained in either culture medium with or without heat inactivated horse serum (2%, Gibco).

### 3. Essays for Viability and Neurite Outgrowth

The viability of cultured neurons was assessed using propidium iodide (PI, Invitrogen). We applied 7.5 μM PI and counted total cell number on brightfield images, while detecting PI+ dead cells with fluorescence, using the Zeiss Cell Observer Spinning Disk (SD) Microscope equipped with a 20x/0.30NA objective. Following the addition of PI, the culture medium was not replaced, and the cells were maintained under standard culture conditions until the subsequent imaging session. Three random regions of interest (ROI) from all plates were imaged every three days until DIV14 (n=4). For the analysis of viability, we subtracted the number of PI+ cells from total cell count and measured the ratio of number of living cells/number of total cells. To quantify neurite outgrowth, we captured brightfield images from multiple ROIs and measured the length of the longest neurites of each cell every three days until DIV14 with a light microscope (Zeiss). Depending on DIV, average 1-45 cells were measured for each time point (n=4). All image analysis for viability and neurite outgrowth were carried out manually using ZEN software.

### 4. Axotomy

To elicit excitation in the cultured neurons to test their responsiveness to injury, *in vitro* axotomy was performed with a UV laser utility (Rapp Opto) integrated on Zeiss LSM880 confocal microscope. The protocol was adapted from [12]. A brief pulse of laser light (337 nm or 355 nm), lasting 0.5-1 second, was directed at axons approximately 50-100 μm from the neuronal soma. The laser emitted approximately 1-30 pulses per second, each with a duration of 3 nanoseconds and an energy output of around 300 μJ. The laser was shot for 1-2 seconds until the cut was visible.

### 5. Calcium imaging

To monitor spontaneous calcium transients in neurons we used Calcium Green™-1 (AM) (Invitrogen, Molecular Probes) staining at DIV30. The medium was replaced with NBA without phenol red containing pluronic acid and Calcium Green™-1 (AM). Cells were incubated for 40 minutes at 21°C, then staining solution was replaced with culture medium. Zeiss Cell Observer Spinning Disk (SD) Time-Lapse Microscope equipped with a 20x/0.30NA objective was used for imaging. Recordings of 1-2-minute periods were taken for detecting spontaneous calcium activity. Calcium imaging combined with axotomy injury was carried out using Zeiss LSM880 confocal microscope as stated above. Calcium activity was observed for one minute as baseline and for one minute after axotomy with time lapse imaging.

### 6. Immunocytochemistry

Cell cultures were fixed on DIV21 in 4% paraformaldehyde (PFA, Sigma) for 15 minutes at room temperature. After PFA was removed, a blocking solution (0.1 mol/L PBS containing 3% Bovine Serum Albumin (BSA), 0.3% Sodium Azide, 0.1% Triton X-10) was added and incubated for 45 minutes at RT. After blocking, the preparation was incubated with primary antibodies in a dilution solution (0.1 mol/L PBS containing 3% Bovine Serum Albumin (BSA), 0.3% Sodium Azide, 1% Tween-20) overnight at 4°C. Beta-III-Tubulin (Abcam-ab107216), Synaptophysin (Santa Cruz-sc-9116), and GFAP (Addgene-114536) were used as primary antibodies. The next day, the samples were washed with PBS and incubated with secondary antibodies (568 goat anti-chicken, 488 goat anti-mouse, and 488 goat anti-rabbit (all from Invitrogen) for 3 hours at room temperature. The preparations were imaged with a confocal microscope LSM800 (Zeiss).

### 7. Statistical Analysis

All statistical analyses were done using Graph Pad Prism 9. Serum and serum free groups were compared using multiple t-test, significance values p < 0.05.

## Results

Firstly, as cloning cylinders were used to facilitate the better attachment of axolotl CNS cells, three different confluency ratios were tested for the area of the cloning cylinders. Cell viability was monitored for 14 days *in vitro* with PI labelling. There were no significant differences detected in terms of viability across serum and serum-free conditions, except at DIV11 and DIV14 in the low confluency group (p value=0.0229 and 0.0065 respectively). Among the three, medium level confluency was found to be the most suitable for other analysis. Cells seeded at other confluency conditions were either too crowded for neurite extension analysis (HC) or too sparse to create a neural network (LC).

The morphological development of the cells *in vitro* was tracked over a period of 14 days. Until the 5^th^ day of culture (DIV5), cells were predominantly round with very little neurite outgrowth. Starting from DIV5, cells exhibited the initial stages of neurite growth (Figure 1C). Cells began to establish connections with neighbouring cells, and a network started to form across the plate by DIV8-11 (Figure 1C). By DIV14, a dense and interconnected network of neurites had formed throughout the culture. The number of neurite-bearing cells in the serum group exhibited a significant increase between DIV5 and DIV8 (p-value = 0.0414); a similar tendency was observed in the serum-free group, albeit not significant (p-value = 0.0707) (Figure 2D). Axon lengths were measured every three days between DIV5 and DIV14. No significant difference in axon length was observed between the serum and serum-free groups during this period (Figure 2E).

**Figure 2.**
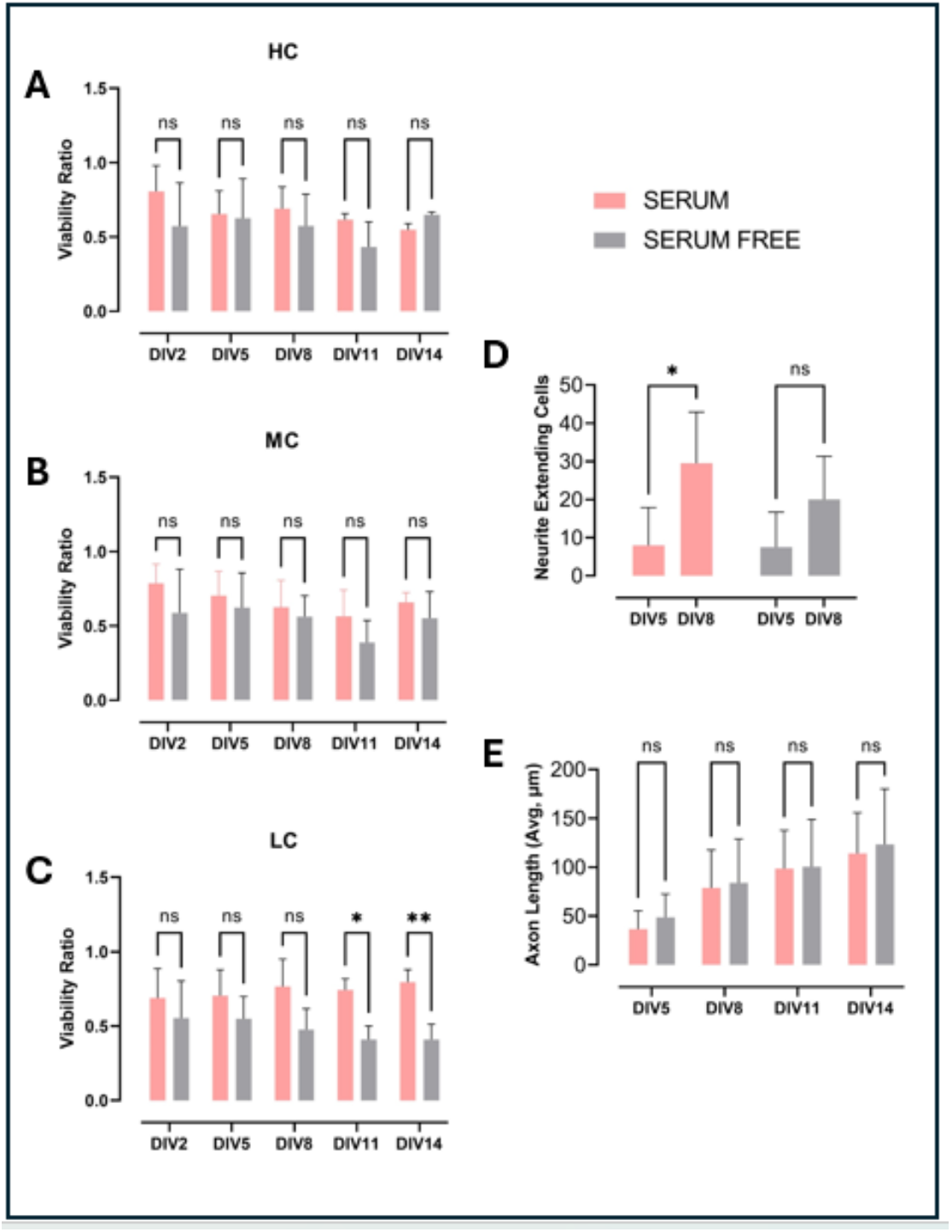
Viability and neurite extension in telencephalic cultures up to DIV14. **A-B-C** Normalized viability ratio of PI negative cells to total cell count in serum or serum-free conditions in high (HC), medium (MC) and low confluency (LC) conditions respectively. **D** Measurement of average maximum axon length over the course of 14 days in serum or serum-free conditions. **E** Neurite extending cell count between DIV5 and DIV8 in serum (p-value = 0.0414) and serum free (p-value = 0.0707) conditions. (n=4)

Spontaneous calcium activity was recorded over a 90-second period using Calcium Green™-1 (AM). The observed spontaneous calcium activity indicates that these cultures contain active neurons with different firing frequencies. The number of spontaneous calcium events varied between 1 to 14 per minute as seen in Figure 3A. We also performed a laser axotomy injury model during calcium imaging to test the responsiveness of the cultures to injury. As seen in Figure 3B and C, axotomized neuron showed significant increase in intracellular calcium level and activity after axotomy.

**Figure 3.**
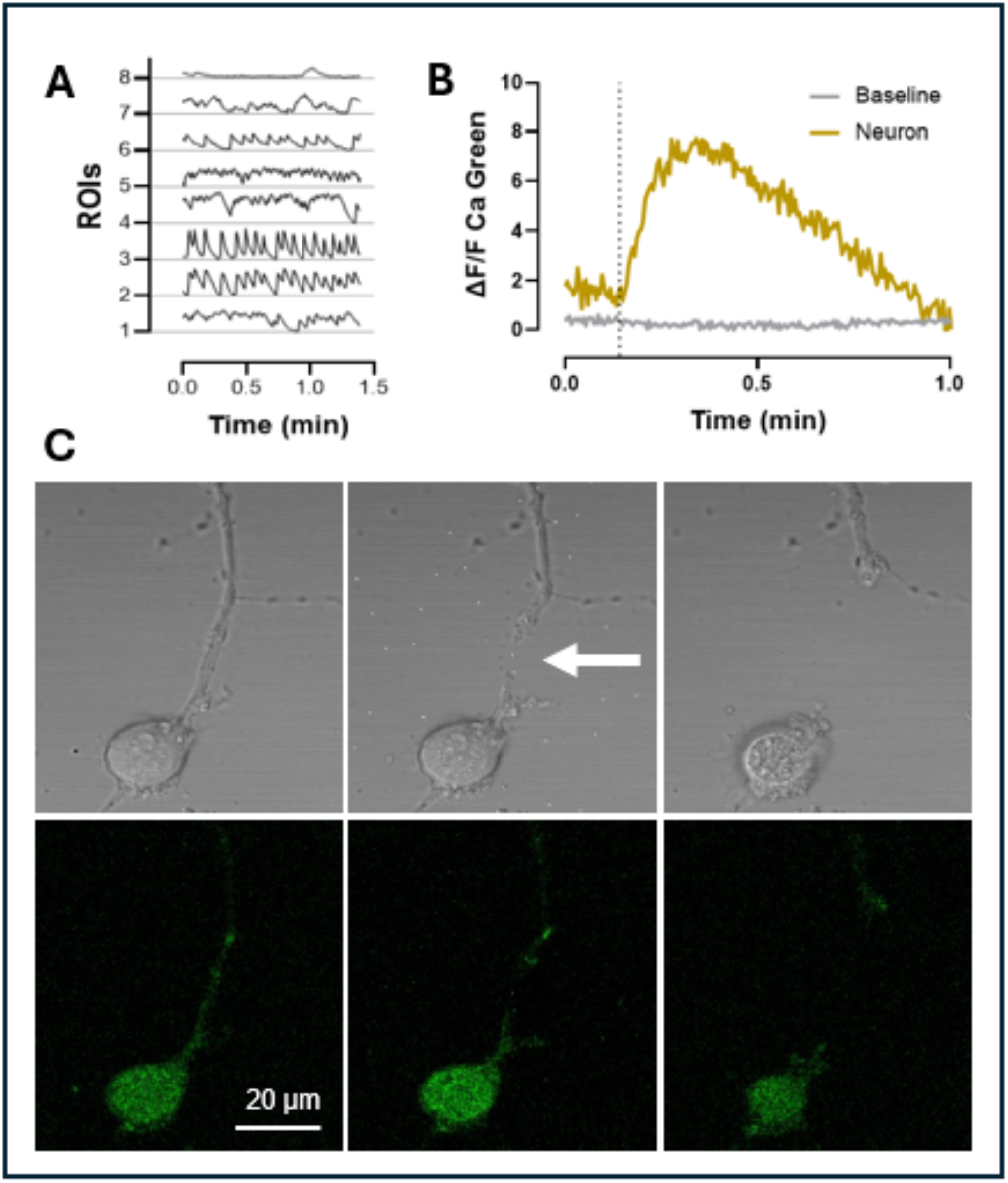
Calcium imaging and axotomy injury **A** Spontaneous calcium activity detected with Calcium Green™-1 (AM) **B** Time series representation of calcium activity in image C, axotomy time point indicated with dashed line. **C** Representative images showing calcium imaging of an axotomized neuron, arrowhead points to the injury site.

On DIV21, we performed immunocytochemistry to identify cell types and morphology of neural network formation. Firstly, βIII-Tubulin and synaptophysin labelling indicated these neurons are capable of making synaptic connections (Figure 4A). While the majority of the cells were identified as βIII-Tubulin+ neurons both morphologically and with neural markers, we also observed some GFAP+ cells. Moreover, we observed βIII-Tubulin immunoreactivity in the nuclei of almost all cell types, which we think was a non-specific cross reactivity (Figure 4B-C) as secondary antibody controls showed no such signal (Supplementary Figure 1).

**Figure 4.**
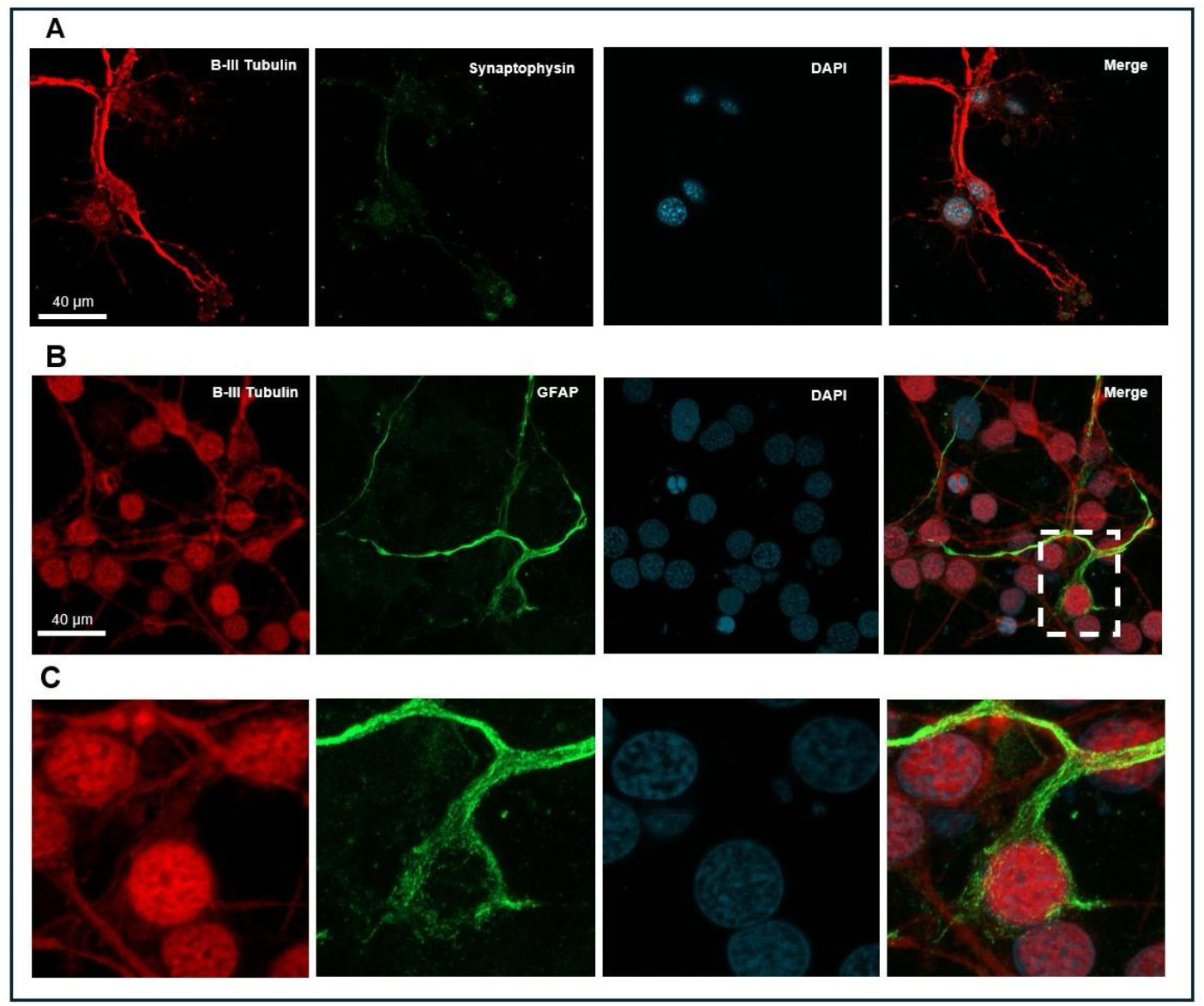
Representative ICC images of axolotl primary telencephalon culture. **A** Cells expressing β-III Tubulin (red), synaptophysin (green) and DAPI (cyan), indicating a neuron identity. **B** Cells stained for β-III Tubulin (red) and GFAP (green) and DAPI (cyan). **C** Inlet image from selected rectangular area showing one cell with immunoreactivity for GFAP (green) in the cytoplasm and for β-III Tubulin (red) in both cytoplasm and the nucleus.

## Discussion

Primary cultures are indispensable to study nerve cells in controlled environment and widely used to investigate physiological and pathological processes of the nervous system. Regeneration of nervous tissue after a damage is an important topic for such studies and often refers to axonal regrowth and less frequently to neurogenesis. Although axolotl is a highly successful organism in both processes there is a lack of an established culture protocol specifically for axolotl nervous system cells. In this study, we successfully developed a method for culturing primary telencephalic cells from the axolotl brain, which yielded both neuron and glial marker positive cells as well as forming a neural network in time.

Several studies demonstrated different methods of culturing various tissues of axolotls; however, there has not been any established cell lines except for the AL-1 line yet. As one of the main focuses of axolotl studies is investigating regeneration, blastema tissue emerging in early stages of limb regeneration is widely studied both *in vivo* and *in vitro*.

There are numerous studies for culturing the blastema tissue [14–16]. There is also one study that demonstrates a protocol for culturing neurospheres from newt brains. They showed neural stem cells (NSC) isolated from newt telencephalon formed neurospheres in culture expressing neuronal stem cell markers [10]. Moreover, after maintenance with differentiation medium with growth factors for 14 days, Tuj1+ neurite extending and GFAP+ cells were detected in these neurospheres [10, 11]. Even though these findings substantiate the highly regenerative capacity of the telencephalic region of amphibian brains, they focus on the presence of stem cell markers and formation of neurospheres in cultures derived from newt brains.

We based the protocol for this axolotl telencephalon culture on our previous primary neuron culture protocol in mice [12] with several adjustments for axolotl cells. In previous examples of amphibian and axolotl primary culture protocols, the temperature and CO_2_ levels for cell maintenance were set more suitable to amphibian physiology. As they are cold blooded animals, cultured cells are better maintained at temperatures between 20-25 °C. Also, CO_2_ levels at 2% were shown to balance pH levels more suitable for these cells [8]. We used L-15 medium for dissection and culturing steps as this medium supports cells in environments without CO_2_ equilibration. We also preferred using NBA medium for culture medium upon seeding the cells as it is designed for long-term maintenance of postnatal and adult brain neurons when supplemented with B-27 [17, 18]. We wanted to test if additional horse serum will make an impact on cell viability. Though, horse serum is known to have a neuroprotective effect on primary neuronal cultures [19, 20]; our findings demonstrate that these cultures can be maintained either in serum or serum-free conditions with slight differences in viability as shown in our results (Figure 2A-E). In limb regeneration studies in axolotls and newts, the complex crosstalk between nerve and blastema tissues was found to involve various growth factors playing a role [21]. Considering aforementioned studies and our results, this finding indicates there is still need for further research on how growth factors influence the regeneration mechanisms in this species. The cloning cylinders we used in the first two weeks upon seeding are commonly used in isolating cell populations in certain culture conditions. However, we observed the conserved area they provide in glass bottom plates also facilitate the attachment of axolotl neurons without clumping when used in combination with PEI coating (Figure 1C), which is usually a challenge in amphibian primary cultures.

We added a centrifuge gradient step to the culture protocol to isolate neuron population from the telencephalon tissue; however, we observed a co-expression of neuron and glia markers in our cultures, with some cells showing both β-III Tubulin and GFAP immunoreactivity. One limitation of immunocytochemistry in axolotls is that there is a lack of reactive antibodies specifically manufactured targeting this species. Therefore, the expression of β-III Tubulin in the nucleus most likely is an artifact but may also indicate differentiating progenitor cells. Secondary antibody controls for Figure 3 showed no such signal in the nucleus (Supplementary Figure 1). In homeostatic conditions, ependymoglia proliferation is observed along the entire ventricular zone in axolotls and increases in injury state [22]. In a recent study, Lust et al. found out that axolotl ependymoglia act as long-term stem cells during homeostatic neurogenesis and can go through a transcriptional change that is specific to injury state [5]. During the dissection and dissociation of neural tissue, the enzymatic and mechanical dissociation in obtaining cultures can also be considered a type of injury to the network of neurons as the neurites get severed [23]. This may be a reason to see increased transcriptional change and, hence, expression of both ependymoglial and neuronal markers simultaneously in our cultures. However, further studies monitoring this expression at longer periods *in vitro* is needed. Our method also allows to model and analyse the neurite outgrowth both at a single neuron and a network level. Neurite outgrowth is an essential feature of neurons in development and also in the adult brain, which makes it an important parameter that is used to monitor and analyse neuronal health and axonal regeneration in *in vitro* studies [24, 25]. Longest neurite outgrowth analysis showed neurons in these cultures can extend over 100 μm average distance in the course of two weeks maintaining the capacity of neuritogenesis and regeneration, which is similar to examples from rat and mice primary neuron cultures in the previously cited studies.

Moreover, the spontaneous calcium activity recorded in our cultures suggests that these neurons retain functional properties, while also showing response to injury induced by laser ablation. This functional aspect is critical for understanding the dynamics of neuronal signalling during regeneration and could provide insights into how axolotls manage to re-establish neuronal networks after injury [4].

Limb regeneration in axolotls is also known to be nerve-dependent both in the initial stages and maintenance [26–28], which makes it crucial to have co-culture models or systems to investigate molecular mechanisms that drive these processes more deeply. This optimised culture protocol provides a valuable tool for future studies aimed at elucidating the regenerative capacity of axolotl CNS neurons and for conducting comparative studies with mammalian neurons to understand the key differences that contribute to their superior regenerative abilities. Furthermore, our findings align with the growing body of literature that emphasises the importance of comparative studies between axolotls and mammals. By understanding the molecular and cellular differences that underpin the regenerative abilities of axolotls, researchers can identify potential targets for enhancing regeneration in less capable species [29]. The insights gained from our axolotl neuronal cultures could thus contribute to broader regenerative biology research, potentially informing approaches to treat neurodegenerative diseases or injuries in humans.

In summary, the development of this culture protocol is the first example of axolotl primary telencephalon culture, and it serves as a platform for exploring the fundamental principles of neurogenesis and regeneration in axolotls. Future studies using this method and also comparisons across species will contribute to unravelling the complex mechanisms that enable axolotls to regenerate their CNS, with implications for regenerative medicine and therapeutic interventions in humans.

**Supplementary Figure 1.**
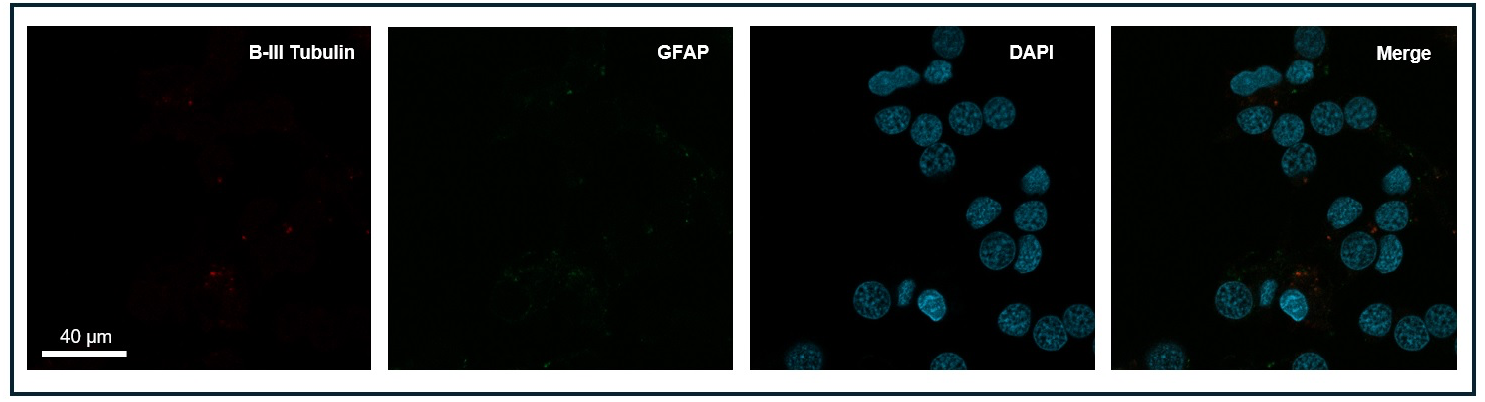
Secondary antibody control ICC images for Figure 3.

